# The function of ethanol in olfactory associative behaviors in *Drosophila melanogaster* larvae

**DOI:** 10.1101/2022.10.13.512092

**Authors:** Michael Berger, Barış Yapıcı, Henrike Scholz

## Abstract

*Drosophila melanogaster* larvae develop on fermenting fruits with increasing ethanol concentrations. To address the relevance of ethanol in the behavioral response of the larvae, we analyzed the function of ethanol in the context of olfactory associative behavior in *Canton S* and *w^1118^* larvae. The motivation of larvae to move toward or out of an ethanol-containing substrate depends on the ethanol concentration and the genotype. Ethanol in the substrate reduces the attraction to odorant cues in the environment. Relatively short repetitive exposures to ethanol, which are comparable in their duration to reinforcer representation in olfactory associative learning and memory paradigms, result in positive or negative association with the paired odorant or indifference to it. The outcome depends on the order in which the reinforcer is presented during training, the genotype and the presence of the reinforcer during the test. Independent of the order of odorant presentation during training, *Canton S* and *w^1118^* larvae do not form a positive or negative association with the odorant when ethanol is not present in the test context. When ethanol is present in the test, *w^1118^* larvae show aversion to an odorant paired with a naturally occurring ethanol concentration of 5%. Our results provide insights into the parameters influencing olfactory associative behaviors using ethanol as a reinforcer in *Drosophila* larvae and indicate that short exposures to ethanol might not uncover the positive rewarding properties of ethanol for developing larvae.

## Introduction

In its natural habitat, the fruit fly *Drosophila melanogaster* prefers to lay its eggs in ethanol-containing fruits [1,2]. Embryos and larvae develop in an environment with steadily increasing ethanol concentrations. The ethanol-enriched environment provides nutrients, increases fitness and protects against predatory insects [3–6]. In the natural environment, ethanol rarely reaches concentrations greater than 5% [7]. Even higher concentrations of ethanol during development can be harmful by increasing malformations in surviving adult flies or by increasing the death rate of developing larvae [1,8].

On the behavioral level, the larvae respond to the presence of ethanol. *Drosophila* larvae move toward a source of ethanol [9]. The attraction to the ethanol odorant does not depend on preexposure and varies slightly at different larval stages and ethanol concentrations [6]. Larvae are attracted to a wide variety of odorants [10], and it is not clear whether the larvae respond with exploratory approach behavior to the presence of an odorant in an odorant- and food-free environment when searching for food or whether larvae associate a positive valence with ethanol.

One way to address whether the larvae perceive ethanol as a positive or negative reward is to associate an environmental cue with ethanol treatment during a training phase and determine after training whether the larvae form a positive or a negative association with the environmental cue. *Drosophila* 3^rd^ instar larvae are a well-established model to dissect the molecular basis of olfactory associative learning and memory [11]. *Drosophila* larvae learn to associate an odorant with a positive taste cue of fructose [12]. Depending on the training cycle, *Drosophila* larvae form shorter- or longer-lasting aversive olfactory associative memories using 2 M NaCl as a negative reinforcer [13]. Training with benzaldehyde or amyl acetate combined with 8% or 20% ethanol resulted in an attraction to the odorant paired with ethanol during the training, suggesting that ethanol can act as a positive reinforcer [6]. In addition to the possible role of ethanol as a reinforcer, a 20-minute exposure to 20% ethanol specifically disrupts the learning of an odorant cue paired with aversive heat shock. This effect is temporary and depends on a non-anesthetic dose of ethanol [14].

Using well-established larval olfactory learning and memory paradigms, we aimed to characterize the role of ethanol as a reinforcer in *D. melanogaster* larvae in more detail. First, we determined whether *Canton S* and *w^1118^* larvae move in or out of substrates containing ethanol and whether ethanol influences odorant attraction. Next, we analyzed whether ethanol functions as a positive or negative reinforcer by varying the training cycles, the reinforced odorant and the order of reinforcement training. Finally, we examined whether the presence of ethanol during the test altered the behavioral outcome. We provide evidence that the concentration of ethanol in the substrate evokes different behavioral responses in larvae depending on the initial presence of ethanol and the concentration of ethanol. The presence of ethanol in the substrate can influence odorant attraction in a concentration- and genotype-dependent manner. The behavioral outcomes in olfactory associative behavior using ethanol as a reinforcer mainly depend on the value the larvae assign to the odorants and the genotype rather than on the presence of ethanol during training. The context of ethanol in the test situation also influences the behavioral outcome.

## Materials and methods

### Fly stocks

Flies were raised on ethanol-free standard cornmeal-molasses food at 25 °C and 60% relative humidity under a 12 h/12 h day/night cycle. *Canton S* and *w^1118^* non-wandering third instar larvae were used for the experiments. To control the growth density of larvae, 35 female flies were mated with 15 male flies and kept for two days in large vials for egg laying.

### Olfactory attraction

To test the larvae for odorant acuity, approximately 20 naïve larvae were transferred to a 2.5% agarose plate containing different concentrations of ethanol (0%, 5%, 8%, 10%). Two odorant cups with perforated lids were set on both sides of the plate. The first cup contained paraffin oil, the solvent for the odorants, and the second cup contained the odorant dissolved in paraffin oil. Odorant balances were determined by placing one cup filled with one odorant on one side of the agarose plate and the second odorant in the second cup on the other side. Larvae were set in the center of the petri dish and covered with a lid. The petri dish was placed under a cardboard box in the hood for five minutes. Next, the number of larvae on each side was counted. In the center of the petri dish, a one centimeter broad stripe was determined to be the neutral zone, where the larvae were treated as undecided. The attraction index (AI) was calculated as follows: number of larvae in the region of odorant A minus the number of larvae in the region of odorant B divided by the total number of larvae (including those in the neutral zone). A negative attraction index reflects an imbalance in favor of odorant B.

### Substrate attraction

To test the larvae for ethanol-containing substrate preference, approximately 20 naïve larvae were transferred to a petri dish containing 2.5% agarose on one side and 2.5% agarose mixed with 5% or 10% ethanol on the other side. Larvae started either on the plain side or on the ethanol-containing side and had 5 minutes to crawl freely in the dark under a cardboard box. Afterward, they were collected and counted, disregarding a neutral zone because a clear border was visible along the middle line of the petri dish. The attraction index was then calculated.

### Olfactory associative behavior

Larval olfactory associative behavior was evaluated based on standardized olfactory training paradigms [12,15]. Briefly, for every experiment, 20 larvae were freshly collected and placed on either a 2.5% agarose plate or an ethanol-containing 2.5% agarose plate with two odorant cups filled with odorant A or B. The plate was covered with a perforated lid, placed in the hood and covered with a cardboard box for five minutes. Next, the larvae were transferred with a brush to a new plate containing the second odorant and again exposed for five minutes. The procedure was repeated one or three times depending on the paradigm used. After training, the larvae were transferred to a test plate with two odorant cups. Depending on the paradigm used, the test plate did or did not contain ethanol. The protocol used to train and test larvae is depicted above the experiments.

### Data analysis and visualization

The box plot summarizes the behavioral data. The top edge of the box plot indicates the 75% quartile, the lower edge is the 25% quartile, and the line within the box is the median. Each whisker reflects 1.5 times the interquartile range. The dots, the edges, the line within the box and the end of the whiskers represent the data values. Dots outside the whiskers are outliers that do not fit within the 1.5 times interquartile range. Significant differences from random choice were determined using the one-sample sign test. Significant differences between two groups were determined with Student’s *t test*, and significant differences among more than two groups were determined with one-way ANOVA with a post hoc Bonferroni Holm correction. Figures were generated with Excel 2016 and GIMP (2.10.12).

## Results

During development, *D. melanogaster* larvae are exposed to increasing concentrations of ethanol in their environment when growing on fruits colonized with yeast [16]. The larvae perceive ethanol as an odorant [6]. Larvae also develop withdrawal-like symptoms after long-term ethanol exposure, suggesting that ethanol elicits rewarding properties [14].

### Higher concentrations of ethanol reduce odorant attraction

Depending on the paradigm used for the olfactory associative learning and memory experiments, during the test, the trained animals had a choice between the two odorants on agarose plates with or without the reinforcer [11]. To analyze whether the presence of ethanol in the substrate influences the motivation of larvae to move, we generated agarose plates with two different areas, where one side contained ethanol and the other did not (Fig 1). We analyzed the behavior of *Canton S* and *w^1118^* larvae after 5 min of exposure to the new environment. The exposure time is identical to the time used to train larvae in olfactory associative learning and memory paradigms (for example, [13]). When *Canton S* larvae were placed on the agarose side, they moved to the border of 5% or 10% ethanol-containing areas (Fig 1A), but the larvae stayed significantly away from the border at 10% ethanol. When they started on the ethanol-containing side, they stayed there independent of the concentration. When *w^1118^* larvae were first placed on the agarose side, they stayed and significantly preferred the ethanol-free side (Fig 1B). When *w^1118^* larvae started on the ethanol-containing side, they stayed in the 5% ethanol-containing side but crawled out of the 10% ethanol-containing side and significantly preferred the ethanol-free area (Fig 1B). It is possible that the larvae stopped at the border of the two areas due to differences in the substrate. To address this, we generated agarose plates with two ethanol-free areas and filmed *Canton S* and *w^1118^* larvae (S1 and S2 Movies). They randomly crawled around and did not aggregate at the border. Thus, the observed behavior of larvae was not due to differences in the surface of the substrate. Taken together, the choice of the larvae to enter or stay on an ethanol-enriched substrate depends on the starting conditions, the ethanol concentration and the genotype.

**Fig 1.**
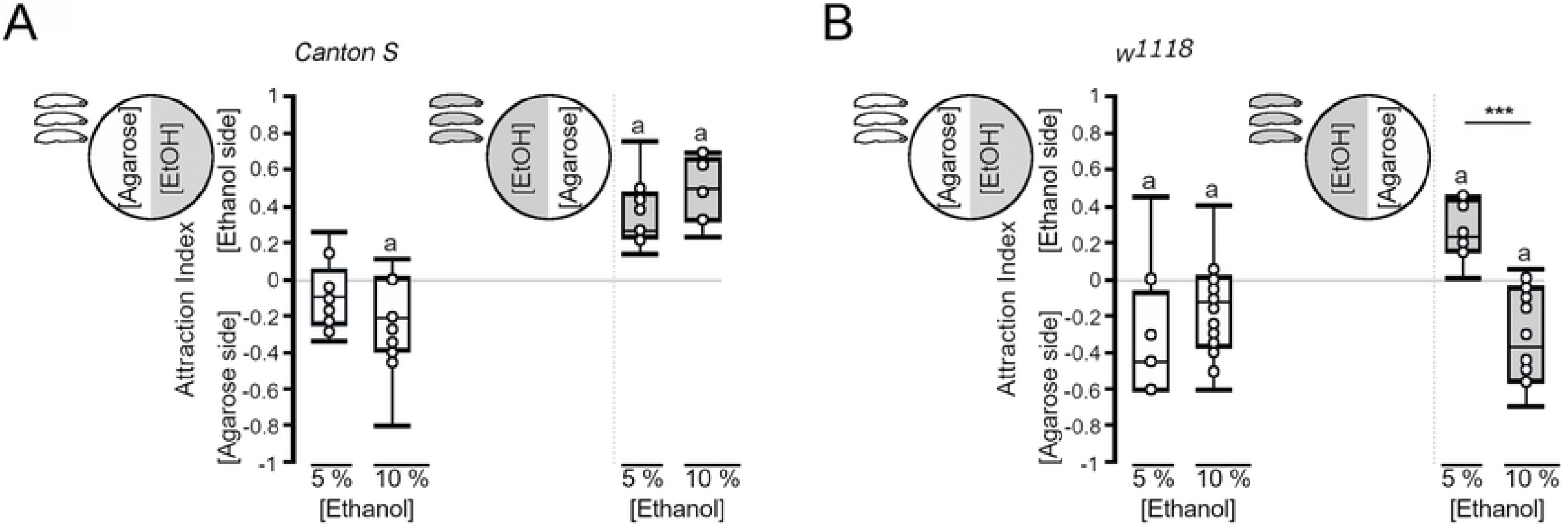
Influence of Ethanol in the Substrate on the Behavior of the Larvae. Placed on agarose, *Canton S* larvae explored the edges of the ethanol-containing area, but they primarily stayed in the agarose area. When starting in an ethanol-containing area, the larvae stayed in the ethanol-containing area (*AI ^agar5%^* = −0.1, ci = −0.22 – −0.05*, AI^agar 10%^* = −0.21, ci −0.38 – −0.04; *AI^EtOH 5%^* = 0.2, *ci* = 0.18–0.38; *AI^EtOH 10%^* = 0.45, *ci* = 0.27−0.59). (B) The *w^1118^* larvae moved to the boundary between the ethanol- and nonethanol-containing areas when they were placed on agarose. When they started on the ethanol-containing area, the larvae stayed in the 5% ethanol-containing area but moved out of the 10% ethanol-containing area (*AI ^agar5%^* = −0.45, ci = −0.6 – – −0.23; AI*I ^agar10%^* = −0.13, ci = −0.34 – −0.01; *AI ^EtOH 5%^* = 0.23, *ci* = 0.15–0.41; *AI ^EtOH 10%^* = −0.38, *ci* = −0.54 – −0.06). N = 8–14 groups of 20 larvae. Significant differences from random choice were determined using the one-sample sign test and are labeled with the letter “**a**”. To determine significant differences between groups, Student’s *t test* was used. *** *P* < 0.001.

To evaluate the significance of ethanol as a positive or negative reinforcer for the behavior of third instar larvae, we wanted to perform associative olfactory learning and memory experiments. In *D. melanogaster* larvae, the odorants amyl acetate and benzaldehyde are commonly used as conditioned stimuli (for example [13]). The odorant 2-heptanone is a naturally occurring component of bananas to which larvae are attracted [17]. First, we analyzed whether ethanol influences the attraction to amyl acetate, benzaldehyde and 2-heptanone in *Canton S* and in *w ^1118^* larvae (Fig 2). Consistent with previous results, *Canton S* and *w ^1118^* larvae were significantly attracted to the odorants amyl acetate, benzaldehyde and 2-heptanone present in odorant cups placed on 2.5% agarose plates (Fig 2A and B). In *Canton S* larvae, the presence of 8% ethanol in the agarose plate significantly reduced the attraction to amyl acetate and 2-heptanone. The attraction to benzaldehyde on 8% ethanol-containing agarose plates was significantly lower than that on 5% ethanol-containing plates but not plates without ethanol. In contrast to *Canton S* larvae, the odorant attraction of *w ^1118^* larvae did not increase or decrease with increasing ethanol concentrations in the substrate. Thus, ethanol reduces odorant attraction in an odorant-specific and concentration-dependent manner depending on the genotype.

**Fig 2.**
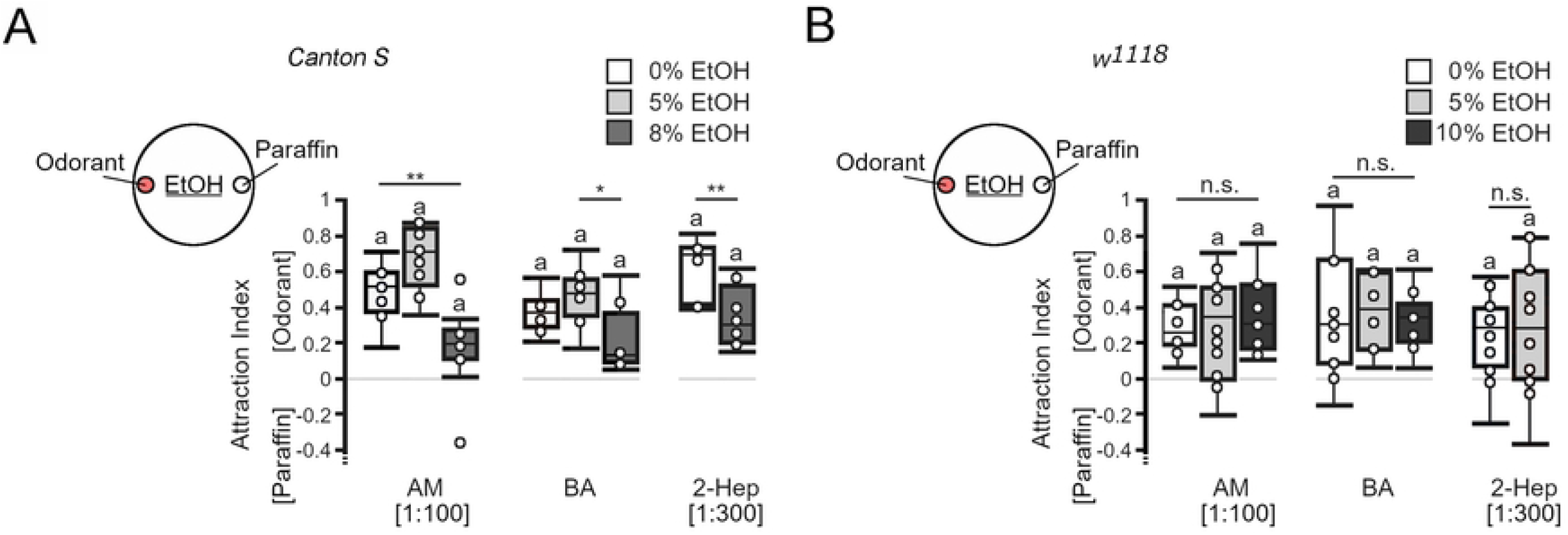
The Effect of Ethanol on Odorant Attraction. The odorant attraction of *Canton S* or *w ^1118^* third instar larvae was determined on 2.5% agarose containing different concentrations of ethanol. (A) *Canton S* larvae showed significantly reduced attraction to amyl acetate (AM) and 2-heptanone (2-Hep) in the presence of 8% ethanol, whereas their attraction to BA differed significantly when larvae were placed on 8% ethanol-containing or 5% ethanol-containing agarose (*AI^AM 0%^* = 0.51, *ci* =0.41–0.59; *AI ^AM 5%^* = 0.7, *ci* = 0.58–0.82; *AI ^AM 8%^* = 0.19, *ci* = 0.11–0.25; *AI ^BA 0%^* = 0.37, *ci* = 0.32–0.43; *AI ^BA 5%^* = 0.48, *ci* = 0.41–0.56; *AI ^BA 8%^* = 0.12, *ci* = 0.1–0.23; *AI ^2-Hep 0%^* = 0.69, *ci* = 0.42–0.72; *AI ^2-Hep 8%^* = 0.3, *ci* = 0.23–0.43). (B) The *w ^1118^* larvae did not change their attraction in the presence of ethanol (*AI ^AM 0%^* = 0.25, *ci* = 0.20–38; *AI ^AM 5%^* = 0.35, *ci* = 0.04–0.49; *AI ^AM 8%^* = 0.31, *ci* = 0.18–0.49; *AI ^BA 0%^* = 0.3, *ci* = 0.09–0.51; *AI ^BA 5%^* = 0.38, *ci* = 0.16–0.58; *AI ^BA 8%^* = 0.33, *ci* = 0.24–0.35; *AI ^2-Hep 0%^* = 0.29, *ci* = 0.12–0.37; *AI ^2-Hep 8%^* = 0.29, *ci* = 0–0.61). N = 8–15 groups of 20 larvae. Two groups were compared using Student’s *t test*, and more than two were compared with one-way ANOVA and a post hoc Bonferroni Holm correction. Significant differences from random choice were determined using the one-sample sign test and are labeled with the letter “**a**”. * *P* < 0.05; ** *P* < 0.01.

### The effect of ethanol during training on odorant choice in an ethanol-free test context

Several requirements must be fulfilled before the effects of ethanol as a reinforcer can be evaluated. First, the odorants used as conditioning stimuli should be perceived by the larvae. We tested this by addressing whether the larvae responded to and moved toward the odorant (Fig 2). Second, the reinforcer should not influence the attraction of the odorant. The presence of 5% ethanol did not significantly change the attraction of *Canton S* and *w ^1118^* larvae to the odorants (Fig 2). Third, to evaluate whether the larvae form an association after training between the reinforced odorant and the nonreinforced odorant, both odorants should be equally attractive and have a similar valence. Initially, *Canton S* larvae did not distinguish between amyl acetate and benzaldehyde (Fig 3A). Over 5 min, the larvae mostly remained close to the middle of the agarose plate, indicating that both odorants were equally attractive. The presence of 5% ethanol in the plate did not interfere with the initial balance (Fig 3A). In contrast, *w ^1118^* larvae showed attraction to amyl acetate on both substrates (Fig 3D). Finally, the training procedure itself should not interfere with the behavior during the test. To test this, we performed one-cycle and three-cycle training with odorants without the reinforcer (Fig 3B, C, E and F). Surprisingly, in these experiments, the *Canton S* larvae were significantly more attracted to amyl acetate than to benzaldehyde when they were trained three times, but the comparison between all groups did not reveal any significant difference due to treatment. The *w ^1118^* larvae showed a significant preference for amyl acetate independent of the treatment. The presence of ethanol in the training plate in combination with benzaldehyde after exposure to an agarose plate with amyl acetate did not significantly alter odorant attractions for *Canton S* or *w ^1118^* larvae (Fig 3B and E). Similarly, pairing amyl acetate with ethanol during training did not significantly shift the balance of *Canton S* larvae (Fig 3C). The attraction to amyl acetate disappeared when *w ^1118^* larvae were trained one time with amyl acetate and ethanol, but the attraction reappeared after three cycles of training (Fig 3F). Overall, there were no significant differences between the different groups. Delaying the test by 5 min after the training did not change the balance of *Canton S* larvae or the attraction to amyl acetate of *w ^1118^* larvae independent of the training condition. Thus, ethanol is not a positive or negative reinforcer for benzaldehyde or amyl acetate in this training and test context for *Canton S* larvae. With one exception in which ethanol represses the attraction to amyl acetate, ethanol did not influence attraction to amyl acetate or aversion to benzaldehyde in the *w ^1118^* larvae.

**Fig 3.**
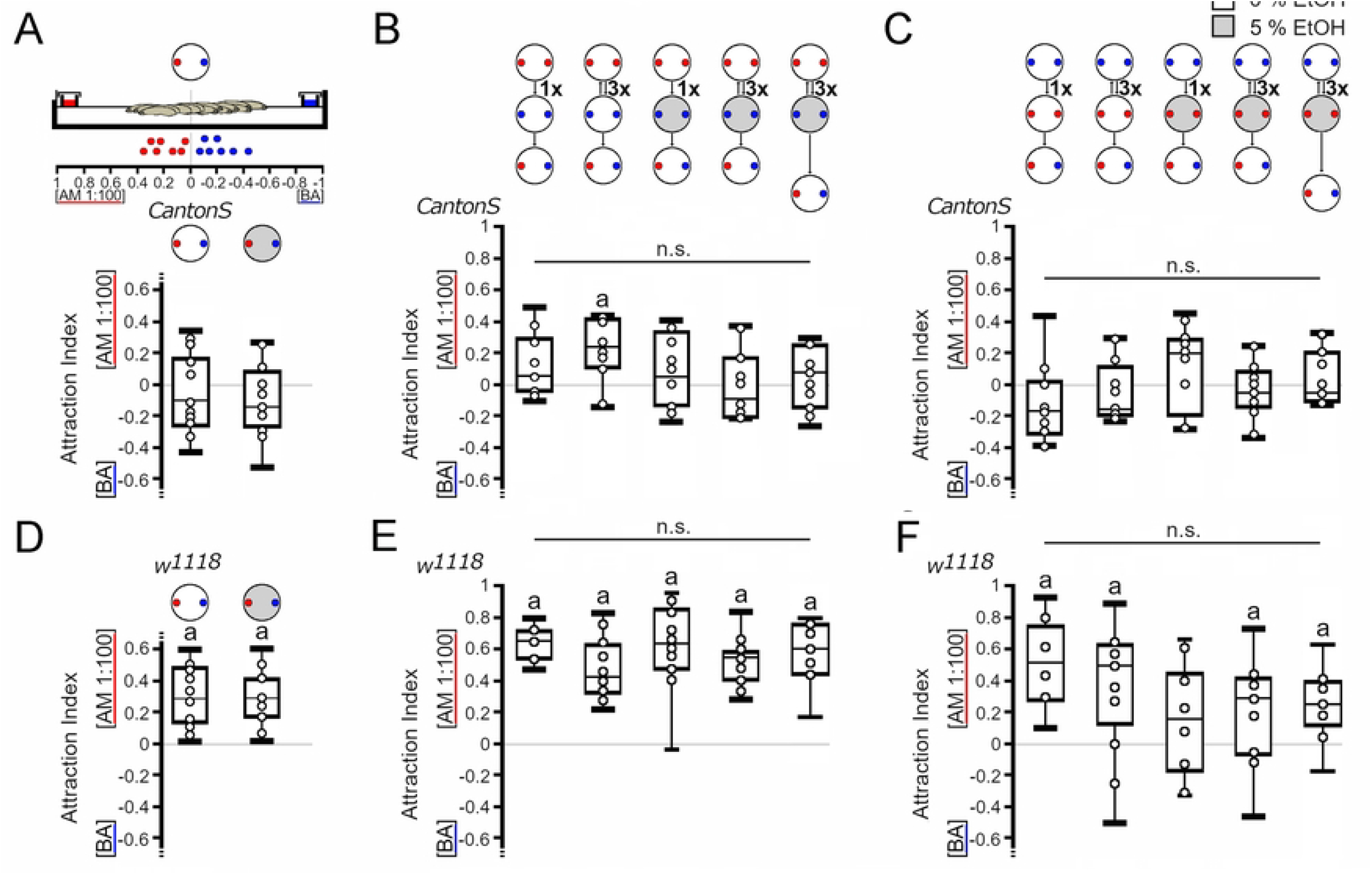
Genotype-Specific Change in Behavior After Odorant Presentation. (A) *Canton S* larvae were equally attracted to both odorants when placed for 5 min on agarose with a source of AM (red dot) and BA (blue dot) (*AI ^balance 0%^* = −0.1, *ci* = −0.22–0.11; *AI ^balance 5%^* = −0.15, *ci* = −0.22–0.03). (B) When larvae were initially exposed to AM followed by BA with the reinforcer, the attraction shifted significantly toward AM after 3 cycles of odorant exposure in *Canton S* larvae. Pairing BA with different concentrations of ethanol did not change the initial balance between AM and BA. The delay of 5 min after training did not change the results (*AI ^AM 1-cycle odor^* = 0.15, *ci* = −0.02–0.23; *AI ^AM 3-cycle odor^* = 0.23, *ci =* 0.10–0.4; *AI ^AM 1-cycle^* = 0.05, *ci* = −0.12– 0.29; *AI ^AM 3-cycle^* = −0.11, *ci* = −0.19–0.13; *AI ^AM 3-cycle+ 5 min^* = 0.1, *ci* = −0.08–0.25). (C) When starting with BA exposure, the larvae were not attracted to AM or BA after one or three cycles of training. Delaying the test for 5 min did not change the balance, and neither did the pairing of AM with ethanol (*AI ^AM 1-cycle odor^* = −0.17, *ci* = −0.28–0.04; *AI ^AM 3-cycle odor^* = −0.16, *ci* = −0.2–0.04; *AI ^AM 1-cycle^* = 0.19, *ci* = −0.07–0.26; *AI ^AM 3-cycle^* = −0.03, *ci* = −0.11–0.09; *AI ^AM 3-cycle+ 5 min^* = −0.06, *ci* = −0.11– 0.14). (D) After 5 min on agarose or 5% ethanol-containing agarose, the *w ^1118^* larvae were significantly more attracted to the odorant (AM 1:100) than to BA (*AI ^balance 0%^* = 0.29, *ci* = 0.15– 0.47; *AI ^balance 5%^* = 0.28, *ci* = 0.17–0.41). (E) When starting the training on agarose in the presence of AM, the attraction of the *w ^1118^* larvae shifted significantly toward AM within 3 cycles of repetition. Pairing BA exposure with different concentrations of ethanol resulted in a significant attraction toward AM (*AI ^AM 1-cycle odor^* = 0.64, *ci* = 0.54–0.7; *AI ^AM 3-cycle odor^* = 0.42, *ci* = 0.33–0.62; *AI^AM 1-cycle^* = 0.63, *ci* = 0.49–0.80; *AI ^AM 3-cycle^* = 0.54, *ci* = 0.44–0.56; *AI ^AM 3-cycle+ 5 min^* = 0.60 *ci* = 0.44–0.75). (F) When the training started with BA exposure, the *w ^1118^* larvae were significantly attracted to AM after one or three cycles of training. Three cycles of training with ethanol resulted in significant attraction to AM. A delay of 5 min before the test did not abolish the attraction (*AI ^AM 1-cycle odor^* = 0.53, *ci* = 0.29–0.66; *AI ^AM 3-cycle odor^* = 0.5, *ci* = 0.26–0.63; *AI ^AM 1-cycle^* = 0.17, *ci* = −0.07– 0.37; *AI ^AM 3-cycle^* = 0.31, *ci* = 0.09–0.44; *AI ^AM 3-cycle+ 5 min^* = −0.25, *ci* = −0.19–0.38). N = 8–17 groups of 20 larvae. Significant differences from random choice were determined using a one-sample sign test and marked with the letter “**a**”, and significant differences between groups were analyzed using one-way ANOVA with a post hoc Bonferroni Holm correction.

### The order of ethanol and odorant presentation might influence the behavioral outcome of the test

To investigate whether the order of the reinforcer representation during training influences the behavioral outcome during the test, we changed the order of training and started with the reinforcement (Fig 4). First, the larvae were trained to associate amyl acetate with 5% ethanol, followed by a transfer to agarose plates with benzaldehyde (Fig 4A and C). They were trained three times. The test was performed immediately after the training and 5 min later. As a control for the procedure, the larvae were only exposed to the odorants during training and then tested. Exposure to odorants without reinforcers resulted in *Canton S* and *w ^1118^* larvae being significantly attracted to amyl acetate. In *Canton S* larvae, amyl acetate combined with ethanol resulted in repression of the attraction; however, when the test was delayed by 5 min, larvae again were attracted to amyl acetate. Since there were no overall significant differences between the groups and the observed repression of attraction was short lived, the phenomenon might be due to the adjustment of the larvae in the test situation rather than being due to associative olfactory short-term memory. Starting the training with benzaldehyde did not shift the balance of *Canton S* larvae. The behavioral outcome differed only in *Canton S* larvae when the order of ethanol-reinforced amyl acetate pairing was exchanged during training (compare Figs 2B and 4A). Here, the delay of the test by 5 min resulted in attraction when the larvae were first trained with amyl acetate and ethanol followed by benzaldehyde exposure but in indifference when the larvae were first exposed to benzaldehyde followed by ethanol-reinforced amyl acetate exposure. In *w ^1118^* larvae, the observed odorant attraction to amyl acetate was not changed regardless of which odorant was first paired with ethanol (Figs 3E and F and 4C and D). Thus, there might be a difference in the behavioral outcome depending on the odorant and the presentation of the reinforcer during training for *Canton S* larvae but not for *w ^1118^* larvae.

**Fig 4.**
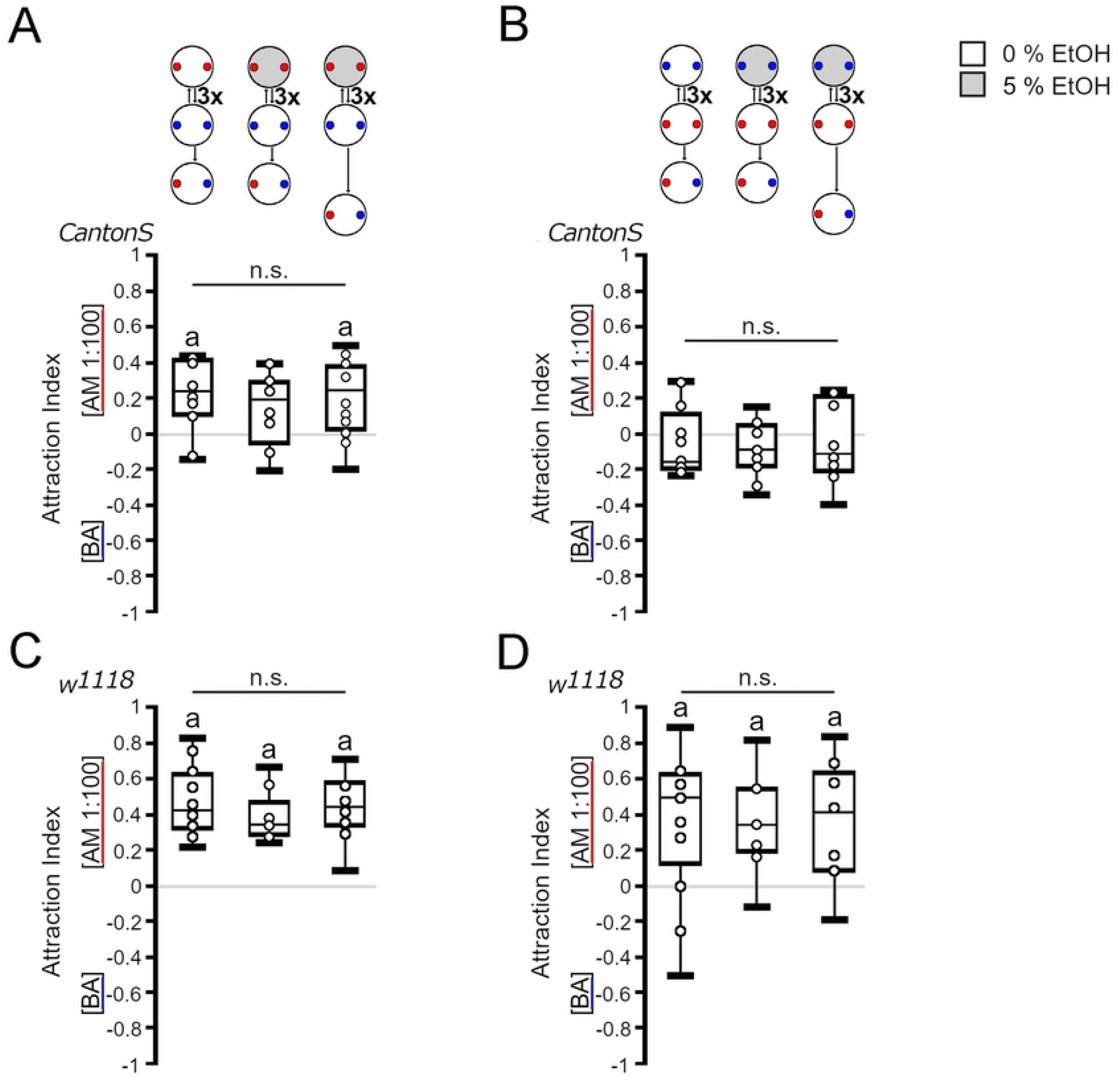
Influence of Order of Reinforcement During Training on Outcome of Test. Odorant attraction after three training cycles starting on ethanol-supplemented agarose. (A) When starting on ethanol-containing agarose with AM exposure, the *Canton S* larvae were not significantly attracted to AM or BA during the test. Extending the training-test interval by 5 min caused a significant attraction to AM (*AI ^AM 3-cycle odor^* = 0.23, *ci* = 0.1–0.4; *AI ^AM 3-cycle^* = 0.19, *ci* = 0.02−0.27; *AI ^AM 3-cycle+ 5 min^* = 0.24, *ci* = 0.05–0.35). (B) When the training started with BA and ethanol, the *Canton S* larvae were indifferent to both odorants in the test. Extending the training-test interval by 5 min did not change the outcome (*AI ^AM odor^* = −0.16 *ci* = −0.2–0.04; *AI ^AM 3-cycle^* = −0.09, *ci* = −0.18–0.01; *AI ^AM 3-cycle+5 min^* = −0.12, *ci* = −0.19–0.18). (C) The *w ^1118^* larvae were significantly attracted to AM in all tested conditions when training started with an AM exposure reinforced with ethanol (*AI ^AA odor^* = 0.42, *ci* = 0.33–0.62; *AI ^AA 3-cycle^* = 0.33, *ci* = 0.29–0.37; *AI ^AM 3-cycle+ 5^ min* = 0.45, *ci* = 0.39–0.58). (D) Similar results were obtained when the training started with BA exposure paired with ethanol (*AI ^AM odor^* = 0.5, *ci* = 0.26−0.63; *AI ^AM 3-cycle^* = 0.33, *ci* = −0.22–0.53; *AI ^AM 3-cycle+ 5 min^* = 0.42, *ci* = 0.11–0.59). (A and C) For comparison, the first boxes show the data from Fig. 3b and 3e. N = 9-12 groups of 20 larvae. The red dots reflect the odorant container with AM, and the blue dots reflect the odorant container with BA. The one-sample sign test was used to determine significant differences from random choice, and the letter “**a**” indicates significant differences. One-way ANOVA with a post hoc Bonferroni Holm correction was used for differences between groups.

To analyze whether the order of the reinforcer presentation during the training influenced the learning index, we calculated the learning index (S1 Fig). To do so, we first calculated the attraction index for each reinforced odorant. Next, we added both attraction indices of the reciprocal groups and divided the sum by two. Depending on which reciprocal group was added to calculate the learning index, *Canton S* larvae perceived 5% ethanol as a positive reinforcer or as a neutral stimulus (S1A and B Fig), whereas *w ^1118^* larvae perceive 5% ethanol as a negative reinforcer or as a neutral stimulus (S1C and D Fig). Delaying the test by 5 min resulted in a random choice between both odorants in the test situation for the *Canton S* larvae. For *w ^1118^*, the delay resulted in indifference or aversion (S1C Fig). When the order of the reinforcer and the odorant were exchanged, *Canton S* and *w ^1118^* larvae did not prefer or avoid the odorant paired with ethanol, suggesting that in this training and test situation, 5% ethanol was perceived as a neutral stimulus.

### Ethanol is a negative reinforcer in a context-dependent manner

To analyze whether the context of the test situation influences the outcome of the training, we next tested the larvae in the presence of the reinforcer after the training cycles (Fig 5). We first addressed whether pairing benzaldehyde with 5% ethanol influenced the choice of *Canton S* and *w ^1118^* larvae on 5% ethanol-containing plates (Fig 5A and D). Next, we performed similar experiments by pairing amyl acetate with 5% ethanol (Fig 5B and E). Depending on the order of reinforcer presentation, the presence of ethanol in the test plate in combination with ethanol-reinforced benzaldehyde resulted in indifference or attraction to amyl acetate in *Canton S* larvae (Fig 5A). In contrast, independent of the order of the reinforcer during training, the pairing of ethanol and amyl acetate resulted in indifference in the test situation (Fig 5B). In *w ^1118^* larvae, training with ethanol-reinforced benzaldehyde (Fig 5C) or amyl acetate (Fig 5D) followed by a test in the presence of ethanol resulted in attraction to amyl acetate with one exception. When larvae were tested in the presence of ethanol directly after training with ethanol and amyl acetate, they did not show a preference for amyl acetate. However, delaying the test by 5 min resulted in attraction to amyl acetate again. To determine the effect of ethanol reinforcement in a test context with ethanol, we calculated the learning index (S2 Fig). For *Canton S* larvae, no clear picture emerged. In some cases, the larvae were indifferent to ethanol, suggesting that ethanol is a neutral cue. In other cases, the larvae formed a negative association with 5% ethanol (S2A and B Fig). In contrast, *w ^1118^* larvae formed a negative association with 5% ethanol in all tested combinations (Fig 2C and D). Thus, 5% ethanol appears to be a negative reinforcer in a context-dependent and genotype-specific manner.

**Fig 5.**
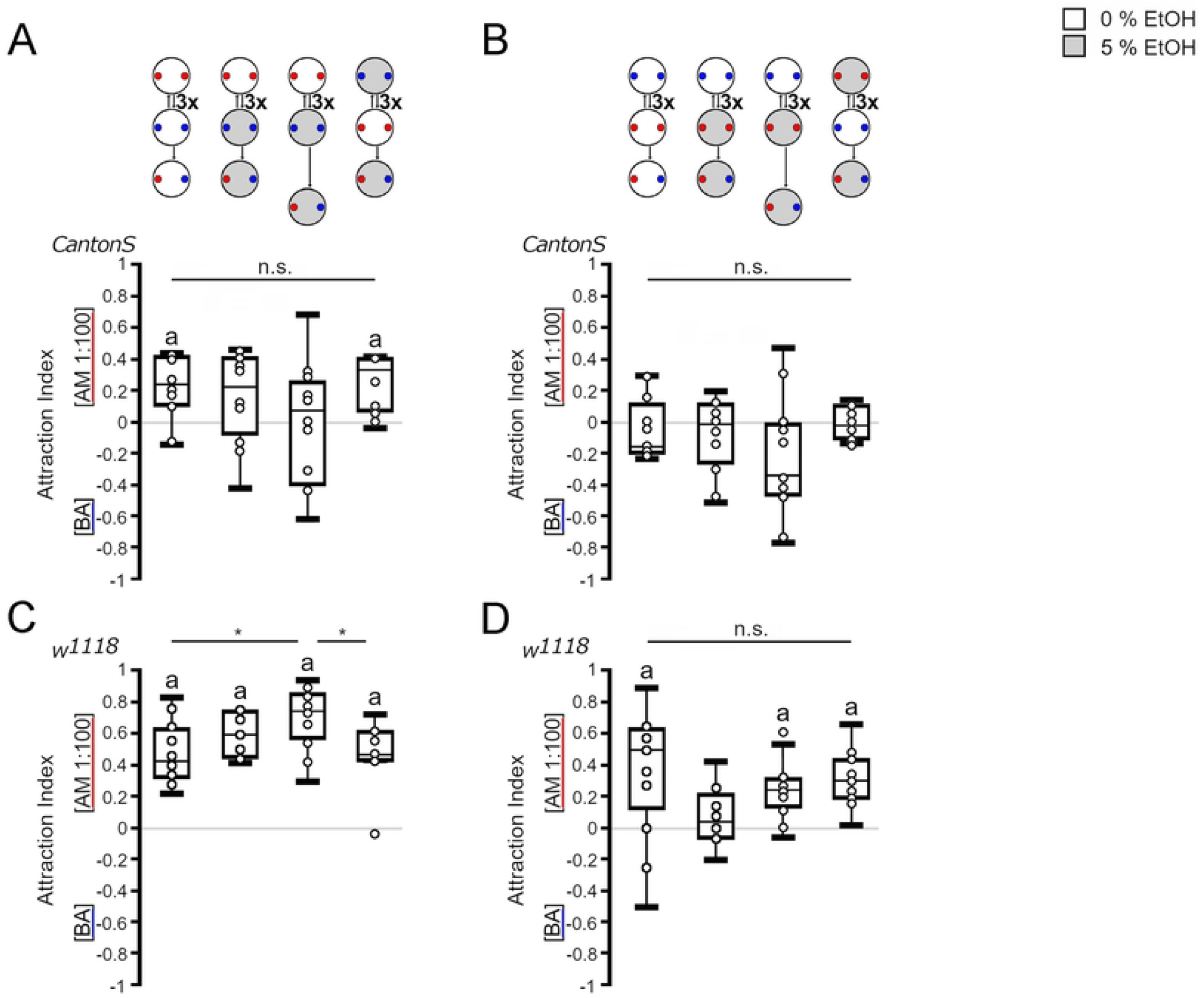
Odorant-Specific Changes in Behavior After Training and Testing with Ethanol. After three cycles of training, odorant attraction was tested on ethanol-containing agarose. (A) *Canton S* larvae were indifferent when the training started with nonreinforced AM but showed attraction to AM when training started with ethanol-reinforced BA exposure (*AI ^AM odor^* = 0.23, *ci* = 0.1–0.4; *AI ^AM 3-cycle^* = 0.22, *ci* = −0.03–0.38; *AI ^AM 3-cycle+5 min^* = −0.07, *ci* = −0.34–0.19; *AI ^AM 3-cycle order^* = 0.33, *ci* = 0.09–0.4). (B) Indifference was also observed when the *Canton S* larvae were first exposed to nonreinforced BA or reinforced AM (*AI ^AM odor^* = −0.16, *ci* = −0.2–0.04; *AI ^AM 3-cycle^* = −0.02, *ci* = −0.18–0.09; *AI ^AM 3-cycle+ 5 min^* = −0.34, *ci* = −0.43–0.04; *AI ^AM 3-cycle order^* = −0.03, *ci* = −0.11– 0.1). (C) When the training started with AM exposure or with ethanol-reinforced BA exposure, the w ^1118^ larvae were significantly attracted to AM (*AIA ^M odor^ =* 0.5, *ci =* 0.26–0.63; *AIA ^M 3-cycle^* = 0.13, *ci* = −0.15–0.31; *AI ^AM 3-c^y ^cle+5 min^* = 0.23, *ci* = 0.17–0.29; *AI ^AM 3-c^y ^cle order^* = 0.29, *ci =* 0.2–0.38). (D) When the training started with BA exposure, the w ^1118^ larvae were indifferent to AM and BA during the test. The delay of the test by 5 min resulted in AM attraction, similar to what was observed when larvae were first exposed to ethanol-reinforced BA during training (*AI ^AM odor^* = 0.5, *ci* = 0.26–0.66; *AI ^AM 3-cycle^* = 0.13, *ci* = −0.15–0.31; *AI ^AM 3-cycle+5 min^* = 0.23, *ci* = 0.17–0.29; *AI ^AM 3-cycle order^* = 0.29, *ci* = 0.2–0.38). N = 9–12 groups of 20 larvae. Significant differences from random choice were analyzed with a one-sample sign test and labeled with the letter “**a**”. More than two groups were compared with one-way ANOVA with a post hoc Bonferroni Holm correction. * *P* < 0.05.

## Discussion

*Canton S* larvae stayed in 5% and 10% ethanol-containing substrates, whereas *w ^1118^* larvae stayed in 5% ethanol but avoided regions containing higher concentrations of 10% ethanol. Staying on ethanol-containing medium can be interpreted as assigning a positive value to ethanol or as indifference to ethanol. In addition, the larvae might not be able to crawl out of the medium due to the sedating and motor impairing effects of ethanol. Since *w ^1118^* larvae moved out of 10% ethanol-containing substrate, it is more likely that *Canton S* larvae assign a different value to ethanol than *w ^1118^* larvae. This is supported by the observation that *Canton S* larvae show a positive association with odorants paired with 5% ethanol in one learning situation as determined by calculating the learning index, whereas *w ^1118^* larvae mostly develop a negative association with odorants with 5% ethanol. A positive association with ethanol has been recently described for olfactory learning and memory experiments for *Canton S* larvae using higher ethanol concentrations of 8% and 20% [6]. However, this raises the question of ecological relevance, since in natural environments, ethanol concentrations are rarely so high [18].

When larvae crawl on ethanol-containing substrates, they come in contact with the substrate, and similar to crawling on fructose-containing agarose plates, the gustatory system is exposed to ethanol. The taste of 10% ethanol appears to be aversive to *w ^1118^* larvae since they crawl out of these ethanol-containing areas (Fig 1B). An aversion to ethanol was also observed in the context of ethanol and food. Female flies lay their eggs in 2% to 6% ethanol-containing food patches [19]. When the larvae hatch, they leave the ethanol-containing food patch [20]. A similar aversion to the taste of ethanol has also been observed in adult flies and honeybees. However, flies and honeybees do not extend their proboscis when the gustatory response is measured using the proboscis extension reflex [21,22].

The valence of ethanol as reinforcer can be evaluated by olfactory associative learning and memory paradigms. In the absence of ethanol during the test, the outcome of the choice between the two odorants in *Canton S* and *w ^1118^* larvae depended on the order of the presentation of the ethanol-reinforced odorant during learning (S1 Fig). *Canton S* larvae form a positive association with 5% ethanol when the reinforcer is first presented with the odorant but not when the reinforcer is presented after exposure to another odorant. The positive association does not last 5 min, suggesting that the response that the larvae show is more likely due to a short-lived shift of odorant valence during training rather than to the positive value of ethanol. This is further supported by the results showing that the shift of valence in the experience of the odorant independent of the reinforcer can also be observed by the attraction to amyl acetate after three cycles of odorant exposure without the reinforcer (Fig 3B). Similarly, in adult flies, exposure to a novel odorant activates specific output neurons of the mushroom bodies that become inactivated when the animal becomes familiar with the odorants [23], indicating that the same odorant is perceived by the brain differently when experienced more often.

The question of whether exposure to odorants combined with 5 min of exposure to 5% ethanol in the substrate is sufficient to trigger the formation of associative olfactory short- or long-term memory thus arises. If accounting for the order of reinforcer presentation during training, *Canton S* and *w ^1118^* larvae do not form a positive or negative association in an ethanol-free test situation. However, *w ^1118^* larvae form a negative association with 5% ethanol when the reinforcer is present in the test situation independent of the order of reinforcement and reinforced odorants (S2 Fig). Normally, after three training cycles using a negative reinforcer of 2.5 M sodium chloride (NaCl), the larvae form a memory that lasts up to 250 min [13]. The observed memory consists of short-term and long-term components [13]. In *w ^1118^* larvae, the observed negative association with 5% ethanol lasted at least for 5 min (S2 Fig), supporting at least a short form of memory that is context dependent.

In summary, the observed positive or negative association with 5% ethanol depends on the order of reinforcement presentation during training, the context of the test situation, the retention interval after training, and the genotype of the larvae.

## Acknowledgments

The authors thank the Scholz lab for fruitful discussions and comments on the paper.

## Supporting information captions

**S1 Fig. Learning Index without Reinforcer During Test.**

The AI values of the learning indices were calculated as follows: [(number of larvae on reinforced odorant side) - (number of larvae on nonreinforced odorant side)]/[(number of larvae on both sides + neutral zone)], unlike the distributional visualization in the previous figures. Each plot is the average of the two reciprocals’ PI values as indicated above the box blot. The data used are from Figs 3 and 4. (A) The learning index of *Canton S* larvae showed that ethanol functioned as a positive reinforcer when the reinforcer was presented at the first odorant exposure, but this effect disappeared when the test was delayed by 5 min (*LI1* = −0.01 *ci* = −0.07–0.08; *LI2* = 0.15, *ci* = 0.07– 0.19; *LI3* = −0.03, *ci* = −0.17–0.05: *LI4* = 0.11, *ci* = 0.04–.0.25). (B) Exchanging the order of reinforcement during training resulted in no significant positive or negative association with 5% ethanol (*LI1* = 0.06, *ci* = 0.03–0.2: *LI2* = 0.05 *ci* = −0.05–.0.12; *LI3* = 0.08 *ci* =-0–0.19; *LI4* = 0.01, *ci* = −0.1–0.2). (C) The learning index of *w ^1118^* larvae showed that ethanol functioned as a negative reinforcer when the reinforcer was presented at the second odorant exposure. This effect did not disappear when the test was delayed by 5 min (*LI1* = −0.11, *ci* =−0.17 – −0.07; *LI2* = 0.025, *ci* = −0.1–0.11; *LI3 =* −0.18, *ci* = −0.28 —0.04: *LI4* = 0, *ci* = −0.11–.0.24). (B) Exchanging the order of reinforcement during training resulted in no significant positive or negative association with 5% ethanol (*LI1* = −0.11, *ci* = −0.12–0: *LI2* = −0.05, *ci* = −0.1–.0.05; *LI3* = −0.08, *ci* =−0.09 – −0.04; *LI4* = −0.04, *ci* = −0.2–0.1). Significant differences from random choice were analyzed with a one-sample sign test and labeled with the letter “**a**”.

**S2 Fig. Learning Index in Test with Reinforcer Context.**

The learning index was calculated as shown in S1 Fig (A) *Canton S* larvae form a negative association when the reinforcer is present at the first odorant exposure (*LI1* = −0.1, *ci =* −0.18 – −0.06: *LI2* = −0.14, *ci* = −0.18 –,−0.07; *LI3* = −0.18, *ci* =−0.22 – −0.09). (B) Changing the order of reinforcement resulted in no association or a negative association of the odorant with ethanol in *Canton S* larvae (*LI1* = −0.09, *ci* = −0.19–0.02: *LI2 =* −0.19, *ci* = −0.23–−.0.1). (C) The *w ^1118^* larvae formed a negative association when the reinforcer was present at the first odorant exposure, and a delay of 5 min after the training did not change the negative association (*LI1* = −0.19, *ci* = −0.38 – −0.17: *LI2* = −0.08, *ci* = −0.15 –.−0.05; *LI3* = −0.28, *ci* =−0.35 – −0.12). (D) Changing the order of reinforcement resulted in a negative association of the odorant with ethanol in *w ^1118^* larvae (*LI1* =−0.18, *ci* = −0.21 – −0.14: *LI2* = −0.21, *ci* = −0.3 –,−0.08).

S1 Movie. Movement of *Canton S* Larvae on 2.5% Agarose with Scarring for 5 min.

S2 Movie. Movement of *w ^1118^* Larvae on 2.5% Agarose with Scarring for 5 min.

## References

1. McKenzie JA, Parsons PA. Alcohol tolerance: an ecological parameter in the relative success of Drosophila melanogaster and Drosophila simulans. Oecologia. 1972;10: 373–388.

2. Richmond RC, Gerking JL. Oviposition site preference in Drosophila. Behav Genet. 1979;9: 233–241.

3. Geer BW, Langevin ML, McKechnie SW. Dietary ethanol and lipid synthesis in Drosophila melanogaster. Biochem Genet. 1985;23: 607–622.

4. Geer BW, Dybas LK, Shanner LJ. Alcohol dehydrogenase and ethanol tolerance at the cellular level in Drosophila melanogaster. J Exp Zool. 1989;250: 22–39.

5. Milan NF, Kacsoh BZ, Schlenke TA. Alcohol consumption as self-medication against blood-borne parasites in the fruit fly. Curr Biol. 2012;22: 488–493.

6. Schumann I, Berger M, Nowag N, Schäfer Y, Saumweber J, Scholz H, et al. Ethanol-guided behavior in Drosophila larvae. Sci Rep. 2021;11: 12307.

7. Gibson JB, Wilks AV. The alcohol dehydrogenase polymorphism of Drosophila melanogaster in relation to environmental ethanol, ethanol tolerance and alcohol dehydrogenase activity. Heredity (Edinb). 1988;60 (Pt 3): 403–414.

8. Ranganathan S, Davis DG, Hood RD. Developmental toxicity of ethanol in Drosophila melanogaster. Teratology. 1987;36: 45–49.

9. Cavener D. Preference for ethanol in Drosophila melanogaster associated with the alcohol dehydrogenase polymorphism. Behav Genet. 1979;9: 359–365.

10. Kreher SA, Mathew D, Kim J, Carlson JR. Translation of sensory input into behavioral output via an olfactory system. Neuron. 2008;59: 110–124.

11. Widmann A, Eichler K, Selcho M, Thum AS, Pauls D. Odor-taste learning in Drosophila larvae. J Insect Physiol. 2018;106: 47–54.

12. Neuser K, Husse J, Stock P, Gerber B. Appetitive olfactory learning in Drosophila larvae: effects of repetition, reward strength, age, gender, assay type and memory span. Anim Behav. 2005;69: 891–898.

13. Widmann A, Artinger M, Biesinger L, Boepple K, Peters C, Schlechter J, et al. Genetic dissection of aversive associative olfactory learning and memory in Drosophila Larvae. PLoS Genet. 2016;12: e1006378.

14. Robinson BG, Khurana S, Pohl JB, Li WK, Ghezzi A, Cady AM, et al. A low concentration of ethanol impairs learning but not motor and sensory behavior in Drosophila larvae. PLoS One. 2012;7: e37394.

15. Gerber B, Hendel T. Outcome expectations drive learned behaviour in larval Drosophila. Proc Biol Sci. 2006;273: 2965–2968.

16. McKenzie JA, McKechnie SW. A comparative study of resource utilization in natural populations of Drosophila melanogaster and D. simulans. Oecologia. 1979;40: 299–309.

17. Oppliger FY, Guerin PM, Vlimant M. Neurophysiological and behavioural evidence for an olfactory function for the dorsal organ and a gustatory one for the terminal organ in Drosophila melanogaster larvae. J Insect Physiol. 2000;46: 135–144.

18. Levey DJ. The evolutionary ecology of ethanol production and alcoholism. Integr Comp Biol. 2004;44: 284–289.

19. Zhu J, Fry JD. Preference for ethanol in feeding and oviposition in temperate and tropical populations of Drosophila melanogaster. Entomol Exp Appl. 2015;155: 64–70.

20. Sumethasorn M, Turner TL. Oviposition preferences for ethanol depend on spatial arrangement and differ dramatically among closely related Drosophila species. Biol Open. 2016;5: 1642–1647.

21. Devineni AV, Heberlein U. Preferential ethanol consumption in Drosophila models features of addiction. Curr Biol. 2009;19: 2126–2132.

22. Mustard JA, Oquita R, Garza P, Stoker A. Honey bees (Apis mellifera) show a preference for the consumption of ethanol. Alcohol Clin Exp Res. 2019;43: 26–35.

23. Hattori D, Aso Y, Swartz KJ, Rubin GM, Abbott LF, Axel R. Representations of novelty and familiarity in a mushroom body compartment. Cell. 2017;169: 956–969.e17.

